# *Wolbachia* successfully replicate in a newly established horn fly, *Haematobia irritans irritans* (L.) (Diptera: Muscidae) cell line

**DOI:** 10.1101/847962

**Authors:** Mukund Madhav, Geoff Brown, Jess A.T. Morgan, Sassan Asgari, Elizabeth A. McGraw, Ulrike G. Munderloh, Timothy J. Kurtti, Peter James

**Affiliations:** Queensland Alliance for Agriculture and Food Innovation (QAAFI), The University of Queensland, Brisbane, QLD 4072, Australia; Department of Agriculture and Fisheries, Brisbane, Australia; Australian Infectious Disease Research Centre, School of Biological Sciences, The University of Queensland, Brisbane, QLD 4072, Australia; Department of Entomology, Center for Infectious Disease Dynamics, Pennsylvania State University, University Park, PA 16802, USA; Department of Entomology, University of Minnesota, USA

**Keywords:** *Wolbachia*, biocontrol, horn fly, *Haematobia* spp., biopesticide, pest management, veterinary ectoparasite

## Abstract

**BACKGROUND:** *Haematobia spp.,* horn flies (HF) and buffalo flies (BF), are economically important ectoparasites of dairy and beef cattle. Control of these flies relies mainly on the treatment of cattle with chemical insecticides. However, the development of resistance to commonly used compounds is compromising the effectiveness of these treatments and alternative methods of control are required. *Wolbachia* are maternally transmitted endosymbiotic bacteria of arthropods that cause various reproductive distortions and fitness effects, making them a potential candidate for use in the biological control of pests.

**RESULTS:** Here we report the successful establishment of a continuous HF cell line (HIE-18) from embryonic cells and its stable transinfection with *Wolbachia* strain *w*AlbB native to mosquitoes, and *w*Mel and *w*MelPop native to *Drosophila melanogaster*. The established HIE-18 cells are typically round and diploid with ten chromosomes (2n = 10) or tetraploid with 20 chromosomes (4n=20) having a doubling time of 67.2 hours. *Wolbachia* density decreased significantly in the HIE-18 cells in the first 48 hours of infection, possibly due to overexpression of antimicrobial peptides through the Imd immune signalling pathway. However, density recovered after this time and stably *Wolbachia*-infected HIE-18 cell lines have now all been subcultured more than 50 times as persistently infected lines.

**CONCLUSION:** The amenability of HF to infection with different strains of *Wolbachia* suggests the potential for use of *Wolbachia* in novel approaches for the control of *Haematobia spp.* Further, the availability of the HIE-18 cell line will provide an important resource for the study of genetics, host-parasite interactions and chemical resistance in *Haematobia* populations.

## 1. INTRODUCTION

*Wolbachia* are maternally inherited endosymbionts of arthropods and filarial nematodes ^1^. They are estimated to infect nearly 40% of terrestrial arthropod species ^2^. The nature of *Wolbachia*-host associations varies from parasitic to mutualistic, often depending on the length of the association ^3, 4^. In longstanding associations, any deleterious effects of *Wolbachia* tend to become attenuated, and *Wolbachia* may eventually form a mutualistic relationship with its host ^1^. Some examples of beneficial effects in such associations are in facilitating parasitoid wasp (*Asobara tabida)* oocyte maturation, female development in adzuki bean borer (*Ostrinia scapulalis)* and the supplementation of vitamin B in bedbugs ^1, 5–7^. The most commonly noted parasitic effects are where *Wolbachia* manipulate the reproductive biology of their hosts to facilitate spread though populations by mechanisms such as cytoplasmic incompatibility (CI), male-killing, feminisation of males and parthenogenesis ^8^. *Wolbachia* may also mediate a range of other effects in their hosts such as the reduction of pathogen transmission, changes in feeding behaviour and locomotion, decrease in offspring survival and reductions in longevity, which can potentially be utilised in the design of programs for the control of arthropods, filarial nematodes and arthropod vectored diseases ^4, 9–13^.

Different *Wolbachia*-host genomic studies have revealed natural horizontal transfer of *Wolbachia* between invertebrate hosts during their evolutionary history ^9, 14^. The mechanism behind horizontal transfer needs to be further explored, but some proposed mechanisms include co-feeding/ salivary exchange, parasitoid wasp or mite co-infection, and faecal-oral transmission ^9, 15–18^. Judging from comparisons of phylogenetic associations, natural horizontal transmission of *Wolbachia* must have occurred relatively frequently in past evolutionary history ^9^. However artificial transinfection is usually more complicated and some species such as red flour beetle *Tribolium castaneum*, silkworm (*Bombyx mori*) and the mosquito *Anopheles gambiae*, appear relatively resistant to *Wolbachia* infection. Despite many attempts it has not yet been possible to develop sustainably transinfected strains of these insects [4]. Also, newly transinfected hosts tend to have a lower infection frequency than in natural associations and often exhibit unstable maternal inheritance ^19–22^. To overcome these difficulties and facilitate the success of cross-species infection, prior adaption of *Wolbachia* to the target host-context by culturing in a cell line from the target host species has been suggested ^9, 21^. For instance, the adaption of *w*MelPop *Wolbachia* (isolated from *Drosophila melanogaster*) in *Aedes aegypti* cells aided the development of a *Wolbachia* infected *Ae. aegypti* mosquito line ^21^. When the mosquito-adapted *Wolbachia* were reintroduced to *Drosophila*, they showed lower virulence, and lower infection frequency in comparison to *Drosophila*-maintained *w*MelPop ^21^. A recent genomic comparison between *w*MelPop grown in *D. melanogaster* cells, a *w*MelPop adapted in mosquito cells (*w*MelPop-CLA) for 3.5 years, and *w*MelPop from *Wolbachia*-infected *Ae. aegypti* mosquitoes (*w*MelPop-PGYP) four years after transinfection, showed genomic differences between *w*MelPop, and *w*MelPop-CLA but no difference in *w*MelPop-PGYP, further indicating that cell line culture can be an excellent medium for quick adaptation of *Wolbachia* to a new host-context ^23^. An additional benefit of culturing *Wolbachia* in cell lines is the ready availability of significant amounts of *Wolbachia* for use in transinfection studies ^4^.

Insects live in diverse ecological niches where they interact with many different species of microorganisms, yet they successfully evade infections because of strong innate immune responses ^24^. Four major pathways (Bacteria: Toll, and Imd; viruses: JAK/STAT and RNAi) are associated with the innate immune system of insects, which is responsible for defense against pathogens ^19, 24–27^. Usually, a Toll signalling cascade is induced by Gram-positive bacteria and fungi, whereas the immune deficiency (Imd) pathway is induced by Gram-negative bacteria ^26, 27^. These signalling cascades increase the production of antimicrobial peptides (AMPs) to suppress or eliminate the infection ^24^.

As the Toll and Imd pathways target bacteria, they can also affect *Wolbachia* density in newly infected hosts ^19^. *Wolbachia*-host interactions are quite variable, and immune genes from Toll (*Cactus*, *Dorsal*, *MyD88*), Imd (*Caspar*, *Relish*, *dFADD*, *IMD*) and AMPs (*Attacin*, *Cecropin*, *Defensin*) are differentially expressed in different hosts ^19, 28–32^. For instance, researchers have found no effect on the innate immune response with the presence or absence of *Wolbachia* in the native host mosquitoes, *Aedes albopictus* and *Aedes fluviatilis* ^19, 28, 33^. However, innate immune genes were overexpressed when *Ae. albopictus* mosquito cells (Aa23) were infected with *w*Mel *Wolbachia*, sand fly cell lines (LL-5) with *w*Mel or *w*MelPop-CLA *Wolbachia*, and *An. gambiae* cell lines with *w*AlbB *Wolbachia* ^19, 34^.

To date, most of these studies have been done in a mosquito or *Drosophila* species context. However, different insect species exhibit different responses to *Wolbachia* ^19^. Clarifying the nature of these interactions in *Haematobia* cells may assist the ultimate development of transinfected strains of *Haematobia* and the design of *Wolbachia*-based *Haematobia* control programs.

Horn flies (*Haematobia irritans irritans*) (HF) and buffalo flies (*Haematobia irritans exigua*) (BF) are closely related obligate hematophagous cattle ectoparasites causing substantial economic losses across the world ^35^. Both species have proven to be highly invasive and they have often been classified as subspecies ^35^. It is hard to morphologically discriminate between them, whereas at the molecular level, ribosomal genes are conserved, and mitochondrial genes have a relatively low divergence of 1.8-1.9% ^36^. The estimated loss associated with the HF alone to North and South America is close to $US3.56B per annum ^35, 37, 38^. Both HF and BF have developed resistance to commonly used chemical insecticides and new methods are required to reduce reliance on chemical controls in endemic areas and to prevent invasion into new areas ^35, 39, 40^.

Here we report the establishment of a HF embryonic cell line (HIE-18) isolated to adapt *Wolbachia* to the *Haematobia* context before BF transinfection. We evaluated the ability of HIE-18 to support growth and development of *w*Mel and *w*MelPop *Wolbachia* isolated from *D. melonogaster* cells, and *w*AlbB *Wolbachia* from mosquito (*Ae. albopictus*) cells. Further, we analysed the host immune response by investigating expression of genes from the Toll and Imd pathways and AMPs to understand the early interactions between *Wolbachia* and the new host HIE-18 cells.

## 2. MATERIALS AND METHODS

### 2.1. Establishment of HF primary cell culture and young cell lines

Embryonated HF eggs were obtained from a laboratory colony reared in the presence of antibiotics and maintained at the USDA/ARS lab in Kerrville, TX, USA^41^. Eggs were sent to the University of Minnesota and processed following the protocol described in Goblirsch et al. (2013) used for the successful establishment of an embryonic cell line from *Apis mellifera* ^42^. Briefly, HF eggs were surface disinfected by sequentially rinsing in a 0.5% sodium hypochlorite and Tween 80 mix, 0.5 % benzalkonium chloride, 70% ethanol and several times in sterile water to remove residual chemicals. The eggs were finally rinsed three times with modified Leibovitz L-15 cell culture medium ^43^ (hereafter referred to as L15C) and gently crushed with a pestle (Kimble Chase, Vineland, NJ) in 1.5 ml sterile Eppendorf tubes to release internal embryonic tissue ^44^. The cellular homogenate was centrifuged at 100 x g for 2 min to remove yolk particles and inoculated in 12.5 cm^2^ non-ventilating screw cap culturing flasks (Falcon, Corning Inc. Tewsbury, MA) in 2 mL of culture medium supplemented with FBS (10%), tryptose phosphate broth (5%), lipoprotein concentrate (0.1%; MP Biomedicals), antibiotics (100 μg/mL penicillin, 100 μg/mL streptomycin, and 0.25 μg/mL amphotericin) (Life Technologies, Grand Island, NY). Flasks were incubated in a non-humidified incubator at 32°C, and media were changed weekly. Confluent primary cultures were dispersed by gentle pipetting and one half of the suspension transferred to a new flask. Young lines (passage 10 or less) were subcultured at a ratio of 1:2 every 2 - 4 weeks and maintained in 25 cm^2^ culture flasks (Cellstar; Greiner Bio-One, Monroe NC) at 32°C. Cell lines in passage 2-5 were frozen by suspending cells in freezing media (L15C supplemented with 20% FBS and 10% DMSO) at the rate of −1°C / min using CoolCell alcohol-free freezing container (BioCision, LLC, Mill Valley, CA) and stored in liquid nitrogen.

### 2.2. Maintenance of *Haematobia irritans* embryonic (HIE) line HIE18

We selected the fastest growing line, HIE18, for continuation. At passage 20, when the line could be split 1:5 every 3-4 weeks, the cells were adapted to Schneider’s medium by gradually increasing the proportion of Schneider’s medium to L15C from 20% to 100% in 20% intervals. Since passage 20, the HIE-18 line has been maintained at 28°C in Schneider’s medium supplemented with 10% FBS.

### 2.3. Other cell lines

*Ae. aegypti* mosquito cell lines (Aag2) infected with *w*Mel and *w*MelPop, provided by the Eliminate Dengue laboratory at Monash University, Australia, and the *Ae. albopictus* mosquito cell line (Aa23T) infected with the *w*AlbB *Wolbachia* strain were cultured in 75 cm^2^ culturing flask with 12 ml of Mitsuhashi and Maramorosch medium (M&M) (Sigma Aldrich, NSW, Australia) and Schneider’s medium (Sigma Aldrich, NSW, Australia) mixed in a ratio of 1:1 at 26°C. The medium was supplemented with 10% fetal bovine serum (Gibco, MD, USA). All cell lines were subcultured once every 6-7 days, and no antibiotics were used during culturing.

### 2.4. Molecular identification of HIE-18 cells

DNA was isolated from whole adult BF and HIE-18 cells using an Isolate II Genomic DNA kit (Bioline, NSW, Australia) following the manufacturer’s protocol. Forward primer HCOI_F 5’-TGAATTAGGACATCCTGGAGCTT-3’ and reverse primer HCOI_R 5’-CACCAGTTAATCCTCCAACTG-3’ were used to target the mitochondrial COI gene of the *Haematobia* species ^45^. Each PCR tube contained 1.5 μl DNA, 1 m of each primer, 1μl of 10x PCR buffer (Qiagen, Melbourne, Australia), 1mM dNTP, 0.1 μl Taq DNA polymerase (Qiagen, Melbourne, Australia) and nuclease-free water to make a total volume of 10 μl. Thermocycling conditions for the reactions were as follows: an initial incubation at 94°C for 3 min, followed by 35 cycles of denaturation at 94°C for 30 sec, annealing at 50°C for 30 sec, and elongation at 72°C for 45 sec with a final extension at 72°C for 10 min. PCR product was amplified in a DNA Engine Thermal Cycler (Bio-Rad, Sydney, Australia) and electrophoresed on a 1% agarose gel. The DNA band was visualised by staining with GelRed (Biotium, NSW, Australia).

### 2.5. Karyotyping

Cell lines in log phase growth were incubated in fresh modified Leibovitz’s L-15 medium containing 12.5 μM colchicine overnight at 32°C. After 12 hours the cells were pelleted at 300 x g for 10 min, resuspended in hypotonic KCl (75mM), and further incubated for 60 min at 34°C. Post incubation HIE-18 cells were fixed in a mixture of methanol and acetic acid (3:1) for 30 min at room temperature. HIE-18 cells were then pelleted and resuspended in fresh fixative. A droplet containing the resuspended cells was placed on a pre-chilled (−20°C) slide to air dry, and stained with 3.2 % Giemsa stain prepared in Sørenson’s buffer (pH=6.8) for an hour at 34°C ^42^. Metaphase chromosomes were counted under an Eclipse E400 phase-contrast microscope (Nikon, NY, USA).

### 2.6. Identification of optimal culturing temperature and media for HIE-18 cells

HIE-18 cells were seeded into 25 cm^2^ flasks with 3.5 ml L-15C at a cell density of approximately 1.1 x 10^6^ cells/ml. Flasks were incubated at 30°C overnight to establish a baseline cell density as a control. The next day flasks were randomly assigned to test temperatures 26-32°C and incubated for six days. On day seven, cells were dislodged by gentle pipetting, pelleted by centrifugation at 300 x g, and incubated in 0.5 ml of NaOH (0.5 N) overnight at 27°C to solubilise cellular protein. The solubilised protein (10 μl) was mixed with 290 μl of Quick Bradford reagent (Sigma Aldrich, NSW, Australia) and incubated at room temperature for 5 min ^42^. Optical density readings for samples were taken at 595 nm using a BMG LABTECH microplate reader (BMG LABTECH, Ortenberg, Germany). Bovine serum albumin (Sigma-Aldrich, NSW, Australia) stock (10mg/ml) in Milli-Q water was used to prepare a protein standard curve. All the experiments were carried out in triplicate. Similar to the above assay, HIE-18 cells were inoculated into 25 cm^2^ flasks at a density of 0.8 x10^6^ cells/ml and incubated overnight to assign a baseline value. The next morning medium from each flask was replaced with one of the test media [modified Leibovitz’s L-15 (Sigma Aldrich, NSW, Australia), Schneider’s (Sigma Aldrich, NSW, Australia), 1:1 mix of Leibovitz’s and Schneider’s, and Shield’s (Sigma Aldrich, NSW, Australia], and cells were left to grow for five days. Protein estimation was carried out according to the previously described protocol.

### 2.7. Transinfecting HIE-18 cells with *Wolbachia*

HIE-18 cell transinfection was carried out using the protocol described by Herbert et al. (2017) ^19^. Briefly, *w*AlbB, *w*Mel, and *w*MelPop infected mosquito cell lines were grown in 75 cm^2^ flasks, each containing 15 ml of M&M media (Sigma Aldrich, NSW, Australia) having 10% FBS. Cells were dislodged from the flask after seven days by vigorous pipetting and spun at 2000 x g. The resulting pellet was washed three times with 5 ml SPG buffer (218 mM sucrose, 3.8 mM KH_2_PO_4_, 7.2 mM MK_2_HPO_4_, 4.9 mM L-glutamate, pH 7.5), and sonicated on ice at 24% power using a Q125 sonicator (Qsonica, CT, USA). Cellular debris was pelleted by spinning at 1000 x g for 10 min at 4°C, and the supernatant was filtered through 50 μm and 2.7 μm acrodisc syringe filters (Sigma Aldrich, NSW, Australia). The filtered supernatant was centrifuged at 12000 x g, and the *Wolbachia* pellet was resuspended in 100 μl of SPG buffer. The pellet suspension was immediately added drop by drop into the cell line flasks when the HIE-18 cells were 80% confluent. The infected cell lines were cultured for 5-6 days and then split in a ratio of 1:2 into the new flasks.

### 2.8. Detection and quantification of *Wolbachia* in HIE-18 cells

DNA was isolated from cells using the Isolate II Genomic DNA kit (Bioline, NSW, Australia) following the manufacturer’s protocol. Detection of *Wolbachia* was carried out by real-time PCR on a Rotor-Gene Q machine (Qiagen, Australia) using strain-specific primers and probes. Primers for *w*AlbB amplified the *wsp* gene (forward primer: 5’-GGTTTTGCTGGTCAAGTA-3’, reverse primer: 5’-GCTGTAAAGAACGTTGATC-3’, probe: 5’-FAM-TGTTAGTTATGATGTAACTCCAGAA-TAMRA-3’) ^46^, whereas the primers used for *w*Mel amplified the *WD0513* gene (forward primer: 5’-CAA ATT GCT CTT GTC CTG TGG-3’,reverse primer: 5’-GGG TGT TAA GCA GAG TTACGG-3’, probe: 5’-CyTGAAATGGAAAAATTGGCGAGGTGTAGG-BHQ-3’) and the primers for *w*MelPop targeted the IS5 element in *WD1310* gene (forward primer: 5’-CTC ATC TTT ACC CCG TAC TAA AAT TTC-3’,reverse primer: 5’-TCT TCC TCA TTA AGA ACC TCT ATC TTG-3’, probe: 5’-Joe-TAG CCT TTT ACT TGT TTC CGG ACA ACCT-TAMRA-3’) ^47^. The total volume of each reaction mix was 10 μl, containing 5 μl of the PrimeTime ® Gene Expression Master Mix (IDT, NSW, Australia), 0.5 μl each of 10 μM forward and reverse primer, 0.25 μl of 5 μM probe and 3 μl of genomic DNA. The optimised amplification conditions for *w*Mel and *w*MelPop were 3 min at 95°C followed by 45 cycles of 10 s at 95°C, 15 s at 51°C, and 15 s at 68°C, whereas for *w*AlbB, amplification was done for 3 min at 95°C followed by 45 cycles of 20 s at 94°C, 20 s at 50°C, and 30 s at 60°C.

For relative quantification of the *Wolbachia* single-copy *wsp* gene (forward primer: 5-’TGG TCC AAT AAG TGA TGA AGA AAC-3’, reverse primer: 5’-AAA AAT TAA ACG CTA CTC CA-3’) and the host *GAPDH* gene (forward primer: 5’-378_F_CCG GTG GAG GCA GGAATGATGT-3’, reverse primer: 5’-445_R CCACCCAAAAGACCGTTGACG-3’) were amplified ^33^. Each sample was run in triplicate on a Rotor-gene Q Instrument (Qiagen, Australia) with total reaction volume of 10 μl containing 5 μl Rotor-Gene SYBR^®^ Green PCR Kit (Bioline, Australia), 0.3 μl each of 10 μM forward and reverse primer and 2 μl of genomic DNA. Negative and positive controls were run with all samples. The optimised thermocycling conditions were initial incubation for 5 min at 95°C followed by 45 cycles of 10 s at 95°C, 15 s at 55°C, and 15 s at 69°C, acquiring green at the end of the step. The relative density of *Wolbachia* was calculated using the delta-delta CT method ^48^.

### 2.9. Effect of *Wolbachia* on population doubling time

*Haematobia* cells infected with *w*AlbB (*w*AlbB-HIE-18), *w*Mel (*w*Mel-HIE-18), *w*MelPop (*w*MelPop-HIE-18) and non-infected cells (HIE-18) were dislodged by pipetting and seeded at a density of 1.5 x 10^6^ cells/mL into three different 25 cm^2^ culture flasks. The flasks were incubated at 28°C in Schneider’s medium (Gibco Life Technologies, Australia) supplemented with 10% FBS. The media were changed every seven days. Cells were visualised under a Motic AE 30 inverted microscope at 40X magnification with a Nikon model DS-L3 camera. A grid (2 cm X 2 cm) was set up on the camera screen (Nikon, Sydney, Australia) for cell counting. Ten randomly selected fields were counted for each flask at 24 h intervals from seeding until they reached confluence >80% ^42^. The experiment was replicated three times, and data for cell density was plotted as the number of cells per cm^2^ (± SD) per day.

### 2.10. Upregulation of immune response genes in *Haematobia* cells

HIE-18 cells were sampled without infection and then at 12, 24, 36 and 48 h post-infection with *w*AlbB, *w*Mel, and *w*MelPop. Total RNA extraction was performed using the Isolate II RNA Mini Kit (Bioline, NSW, Australia) following the manufacturer’s protocol. Primers were designed as detailed below to target the *H. irritans* innate immune genes *cactus*, *relish*, *attacin*, *cecropin*, and *defensin*. Targeted gene sequences of *D. melanogaster* were collected from the Flybase web server (https://flybase.org/) and searched on the NCBI database to find similar sequences in *Musca domestica*, the closest species to *Haematobia* for which gene sequences were available. Gene sequences from *M. domestica* were searched for sequence similarity in the recently sequenced *H. irritans* genome (Accession number: GCA_003123925.1) to find similar contigs. Finally, primers were designed based on those contigs using Genious Prime software (See Table 1). A SensiFAST Probe No-ROX One-Step Kit (Bioline, NSW, Australia) was used to carry out the gene expression study following the manufacture’s protocol with a Rotor-gene Q Instrument (Qiagen, Australia). Melt curve analyses were performed in the range 55– 90°C to examine reaction specificity of the designed primers. All of the assays were performed in triplicate. Gene expression was normalised to the host *GAPDH* gene, and change in expression level was calculated using the delta-delta CT method ^48^.

**Table. 1.**
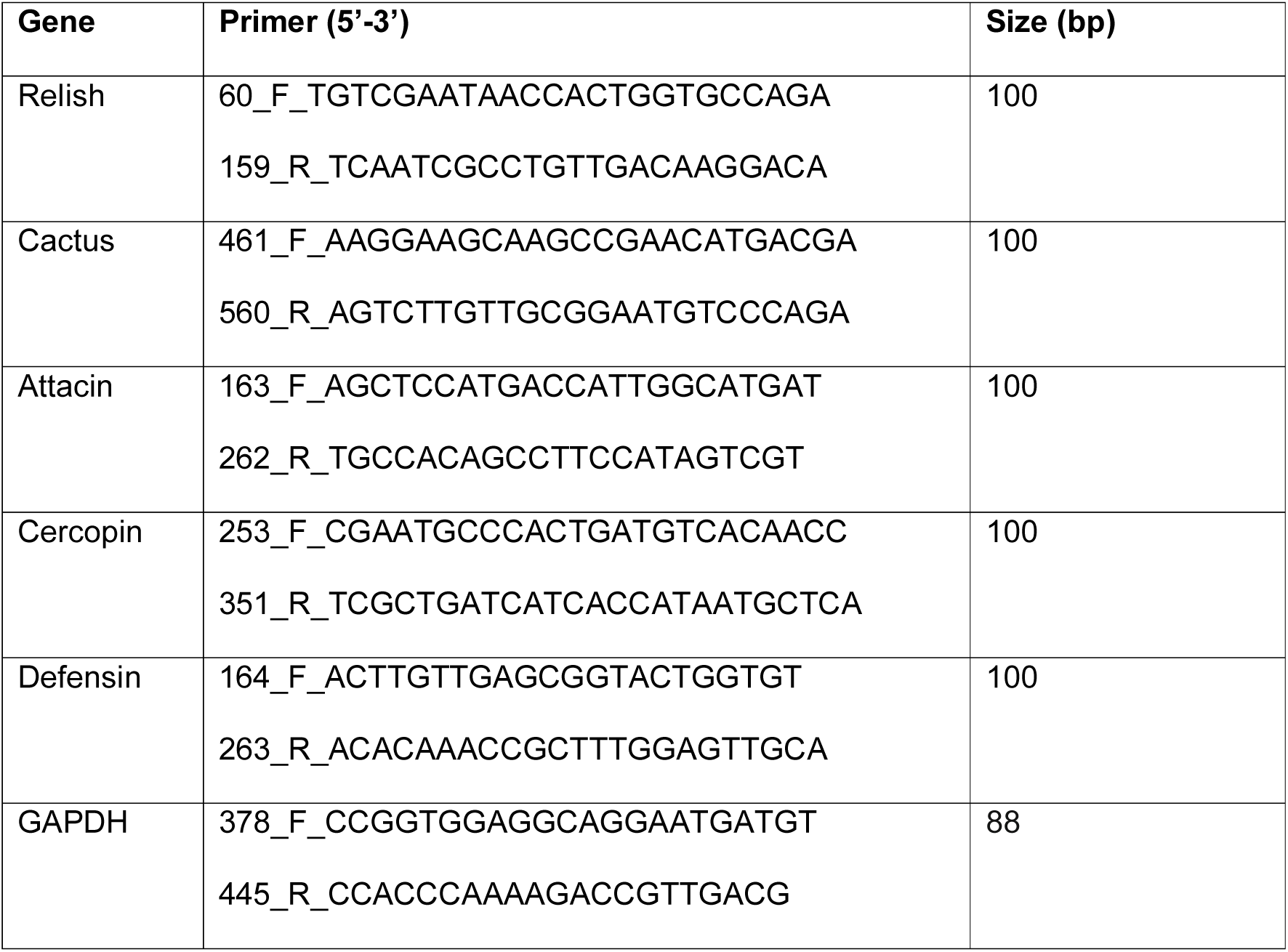
List of primers designed to study immune response to infection with *Wolbachia* in *H. irritans*.

## 3. RESULTS

### 3.1. Features of HIE-18 cells isolated from HF embryos

Twenty-two primary cultures were made to initiate primary cell lines from HF eggs. In some instances, a slight variation was made to cell establishment protocols, either by changing the seeding quantity of the inoculant (50 - 200 embryos) or centrifuging cellular homogenate to remove the excess of yolk material. We expected that having more cells from embryos would help in increasing cell-cell interaction, and extra yolk material would provide the necessary nutritional support for cellular growth. However, an excess of yolk material was found to interfere with cellular adherence in the flasks. One culture of HIE-18 cells from HF embryos grew in aggregates of 20-30 cells where cellular morphology was hard to discriminate in the first two-three weeks (Fig. 1A). Nevertheless, within eight weeks cells reached confluence and a distinct morphologically heterogeneous population of round epitheliocytes, neuronal-, and fibroblast-like cells were seen. Later, round cells (5-10 µm) became predominant. These were isolated and cryopreserved (Fig. 1B). Currently, HIE-18 cells have been successfully passaged over 200 times (Fig. 1C). Ten of the remaining 22 primary cultures developed into lines (passage 7 or less) and are currently stored in liquid nitrogen.

**Fig. 1.**
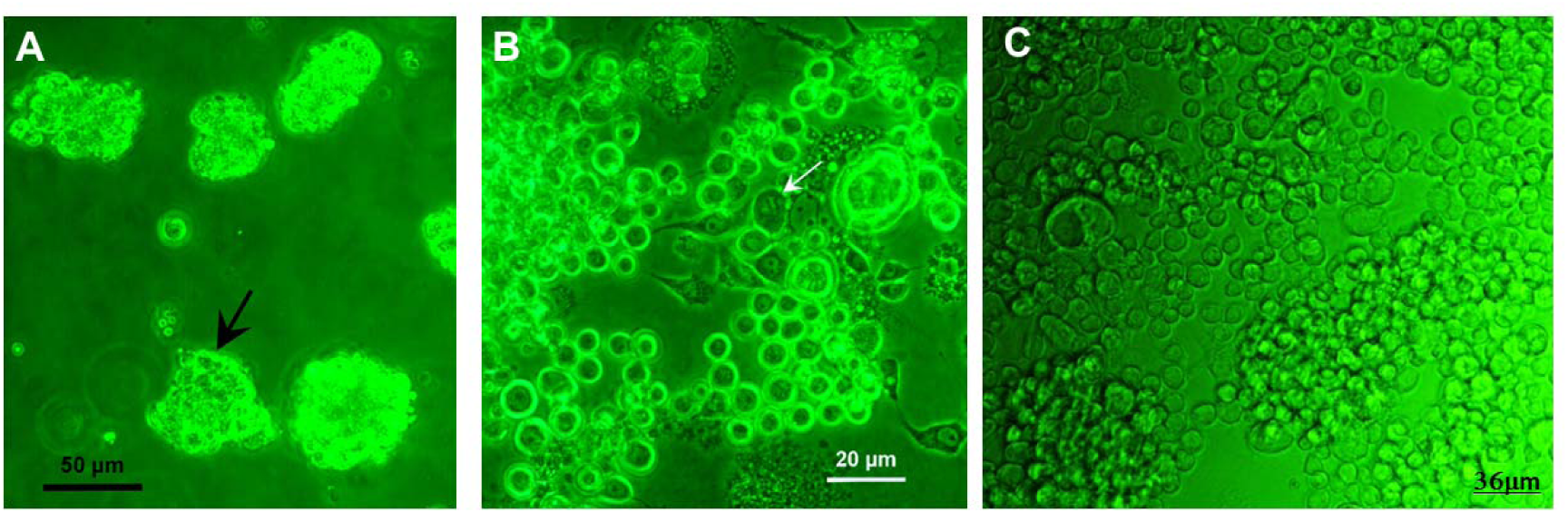
Cellular morphology and features of *H. irritans* cell cultures. **A**. Embryonic fragments seeded into primary cultures, one-day post-inoculation; arrow indicates a cluster of 20-30 cells. **B**. Round cells predominate in young HIE18 cultures, third transfer, two months in culture. Arrow indicates a cell in metaphase. **C.** Current continuous HIE-18 isolate cells (June 5, 2019) maintained in Schneider’s medium. The cells are predominantly round, 5 - 10 µm, loosely adherent and form multicellular clusters.

### 3.2. Species confirmation using PCR and chromosomal analysis

HIE-18 cell identity was confirmed by amplifying the *COI* gene of approximately 920 bp size with *Haematobia*-specific primers. Both the HIE-18 cells and positive control BF were amplified (Fig. 2A). HIE-18 cells were comprised predominantly of round diploid cells with ten mitotic chromosomes or tetraploid cells with 20 chromosomes (Fig. 2B-C). Diploid cells had five pairs of chromosomes, two of which were sub-metacentric and three metacentric. There were no heteromorphic (sex) chromosomes.

**Fig. 2.**
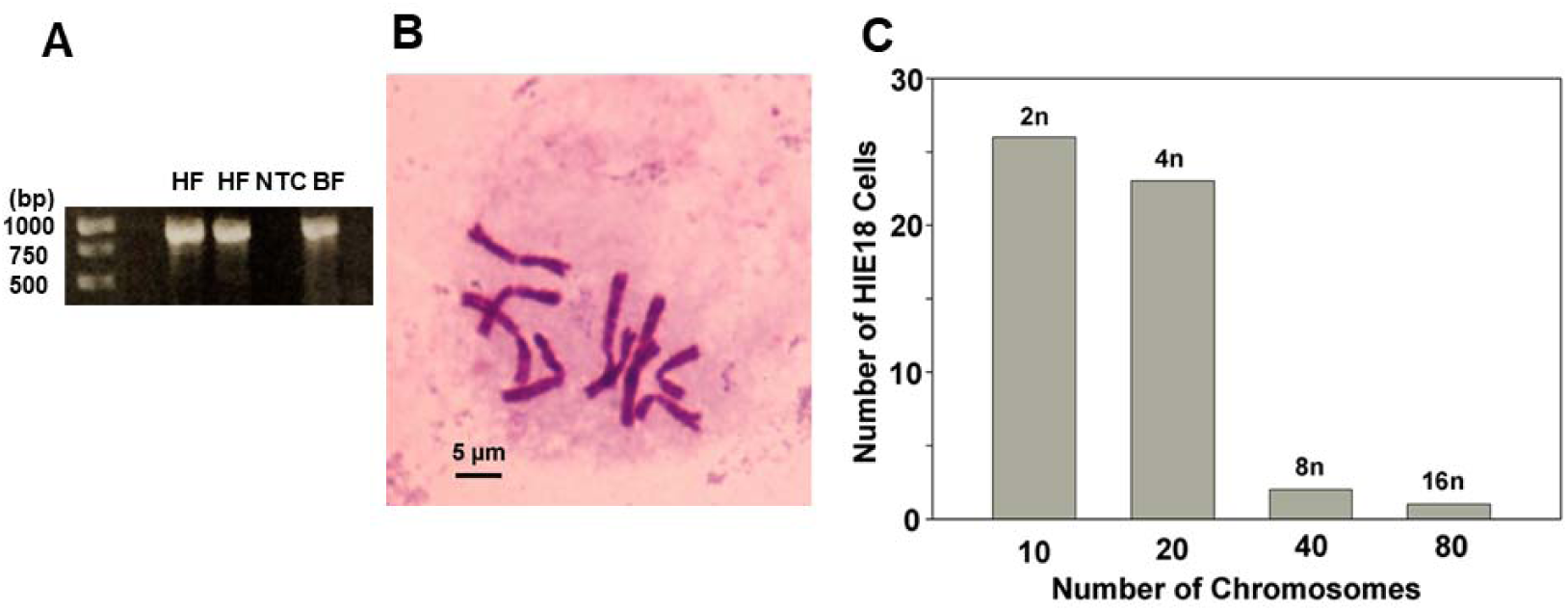
PCR and chromosome profile confirmation of HIE18. **A.** Approximate 920-bp product amplified using mitochondrial-encoded COI primer specific to *Haematobia* spp. HF: *H. irritans* HIE-18 cell line, NTC: Negative control, BF: *H. exigua*. **B.** Metaphase chromosome spread in Giemsa stained diploid HIE-18 cells. **C.** Graph shows a number of metaphase chromosomes in *H. irritans* cells.

### 3.3. Identification of optimal culturing temperature and media for HIE-18 cells

Protein concentration was used as a measure of growth of the newly established HIE-18 cell line incubated at four different temperatures (26, 28, 30 and 32°C) (Fig. 3A). There was a significant difference in cellular replication at different temperatures (one-way ANOVA: *F*_4,10_ = 237.1, *p*<0.0001). Highest protein concentration was seen at 30°C (2.6+0.72 mg), followed by 28°C (2.33+0.79 mg) indicating the best growth at these two temperatures. There was also a significant difference in the concentration of protein in different culturing media (one-way ANOVA: *F*_4,10_=118.3, *p*<0.0001) (Fig. 3B) with the HIE-18 cells growing best in Schneider’s medium (2.74 + 0.11 mg) and 1:1 mix of Schneider’s and L-15 culture media (2.49 + 0.13 mg).

**Fig. 3.**
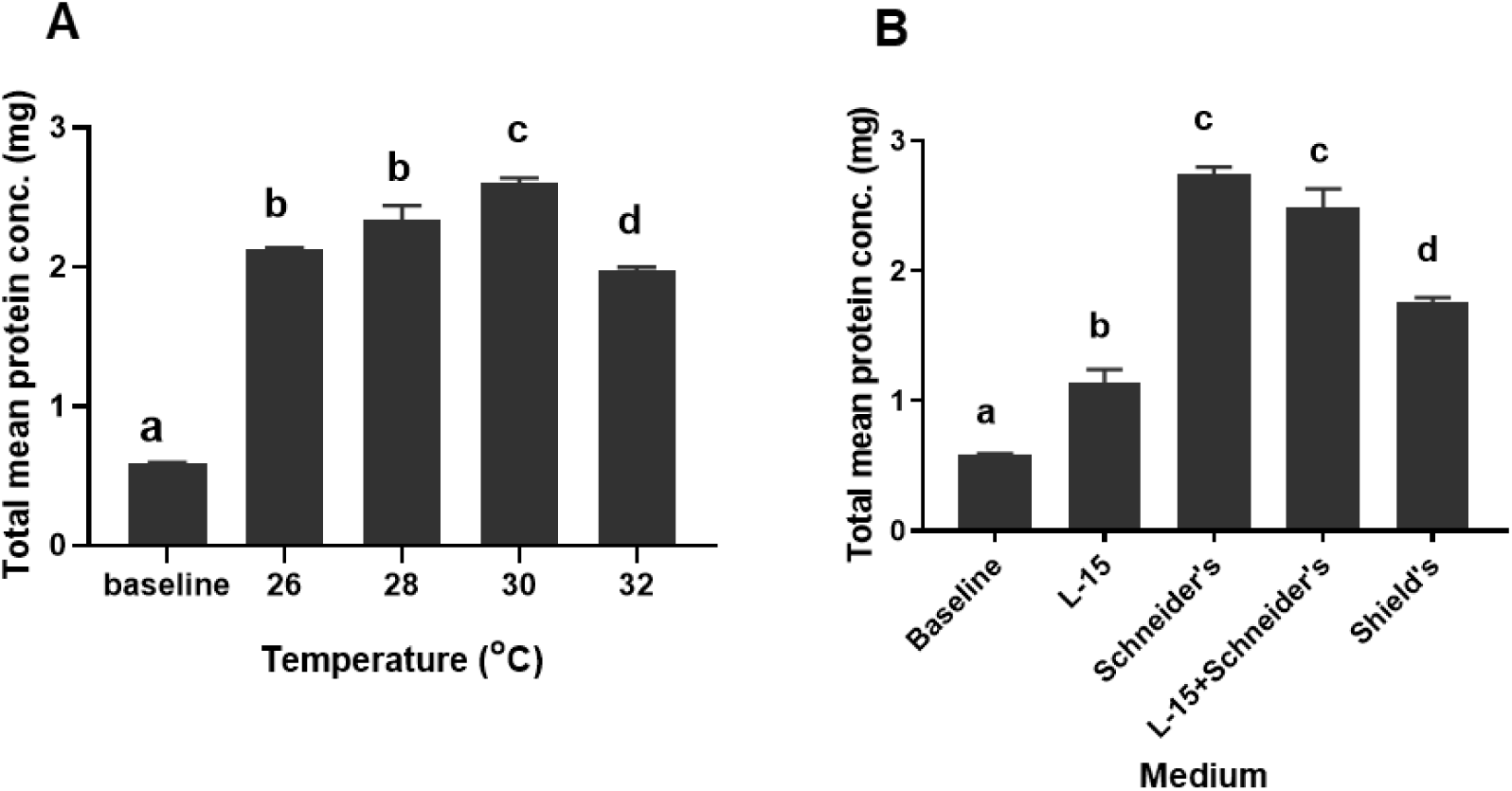
Protein production in HIE-18 cells grown in different temperatures and culturing mediums. Left axis shows protein concentration used as a proxy for the HIE-18 cellular growth, determined using a Bradford assay. Here, baseline data is protein production from the HIE-18 cells grown in modified L-15 medium for 24 hours at 30°C before assigning test temperature and test medium. **A.** Total protein production was significantly different between the cultures incubated at different temperatures (*F* _4, 10_= 237.1, *p*<0.0001). **B.** Total protein production was significantly different between the cultures grown in different culturing media (*F* _4, 10_= 118.3, *p*<0.0001). Error bars are SEM calculated from three replicates of culture flasks. Columns with different letters are significantly different (Tukey’s multiple comparison test; *p* = 0.05).

### 3.4. *Wolbachia* replication in HIE-18 cell lines

It took multiple rounds of *Wolbachia* introduction into the HIE-18 cell lines to establish persistent infections of *w*AlbB (two rounds), *w*Mel (three rounds) and *w*MelPop (two rounds). We saw a significant decrease in the density of *Wolbachia* within 48 hours of infection: *w*AlbB (one-way ANOVA: *F*_3, 8_ = 10.87, *p*=0.0034), *w*Mel (one-way ANOVA: *F*_3, 8_ = 27.30, *p*=0.0001), and *w*MelPop (one-way ANOVA: *F*_3, 8_ = 56.34, *p*<0.0001; Fig. 4A-C). To date, HIE-18 cells infected with *w*AlbB, *w*Mel, and *w*MelPop have been successfully maintained for 70, 60 and 50 passages, respectively, with stable infection and no significant change in *Wolbachia* density over these passages (Fig. 4D-F). Interestingly, *w*AlbB appeared to grow at lower density than the other two strains, *w*Mel and *w*MelPop, in HIE-18 cells (Fig. 4A-F).

**Fig. 4.**
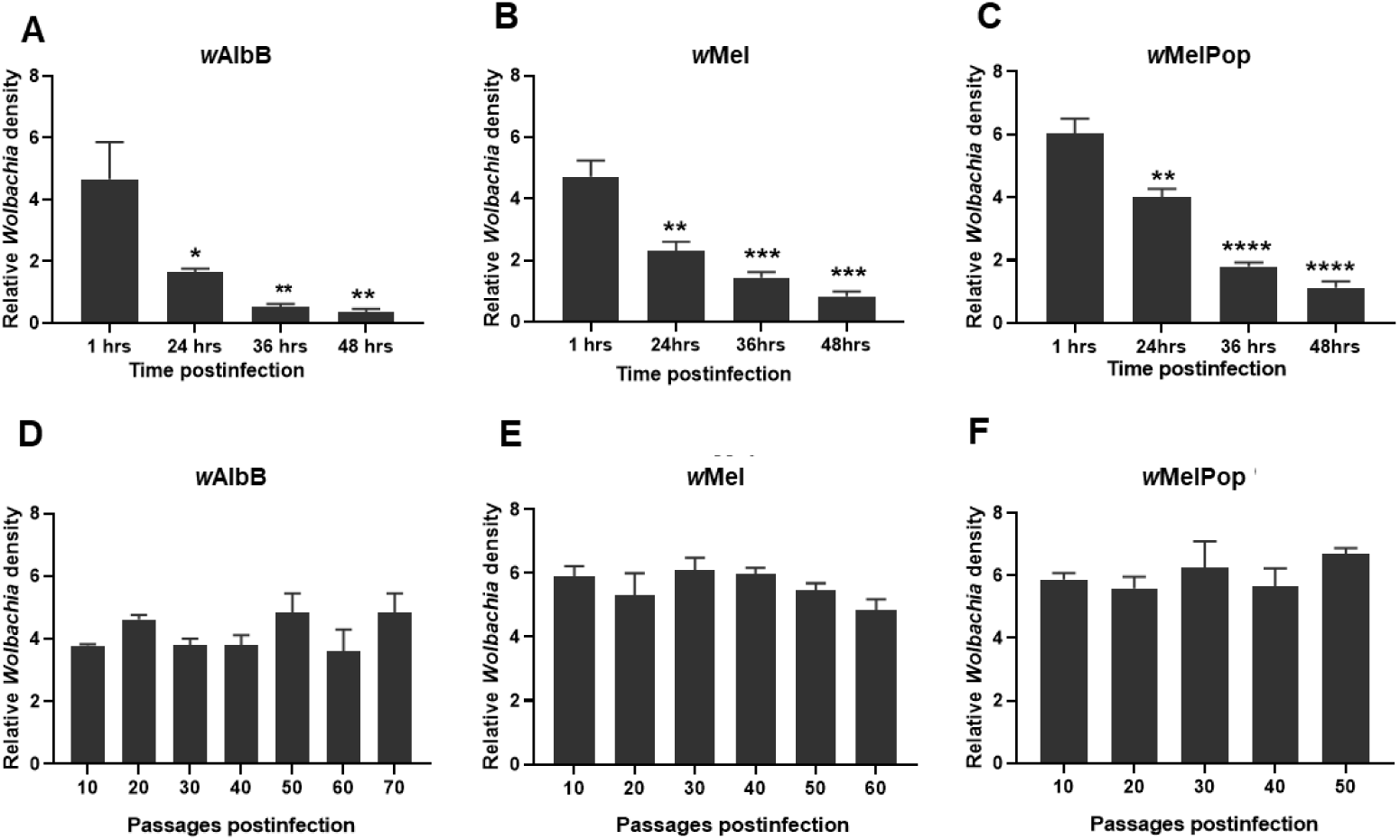
Quantification of *Wolbachia* (*wsp*) relative to host (*GAPDH*) using real-time PCR in HIE-18 cell post-infection. **(A-C).** Significant decreases in *w*AlbB, *w*Mel and *w*MelPop were observed within 48 hours post-infection. Bars with asterisks represent significant difference compared to one hour post-infection using Dunn’s test; \**p*<0.02, \*\**p*<0.003, \*\*\**p*<0.0003, \*\*\*\**p*<0.0001. **D.** *w*AlbB dynamics in HIE-18 cell line over 70 passages. **E.** *w*Mel dynamics in HIE-18 cell line over 60 passages. **F.** *w*MelPop strain dynamics in HIE-18 cell line over 50 passages. Tukey’s test (*p*= 0.05) showed no significant variation in *Wolbachia* infection over passages (D-E). Error bars indicate SEM from triplicate culturing flasks.

### 3.5. Effect of *Wolbachia* infection on the population doubling time of HIE-18 cells

Population doubling time was calculated after stably infecting HIE-18 with *w*AlbB, *w*Mel, and *w*MelPop strains. The population doubling time for the uninfected HIE-18 cells was approximately 67.0 hours and the slope of the cell number by timeline plot consequently higher than the infected lines (y = 7.2032x + 12.088; r^2^ = 0.998) (Fig. 5). For the *w*Mel-HIE-18 (77.0 hours) and *w*AlbB-HIE-18 (79.2 hours) doubling time was longer and the slope of the population growth lines consequently not as steep (6.479x + 11.716; r^2^ = 0.997; 5.7843x+11.756, r^2^ = 0.994). The *w*MelPop-HIE-18 grew more slowly than the others with a doubling time of approximately 86.4 hours (y = 4.363x + 13.036; r^2^ = 0.992). The newly introduced *Wolbachia* infections appear to exert varying levels of stress on the new host cells, increasing their doubling time and this effect was most marked in the case of the *w*MelPop infection.

**Fig. 5.**
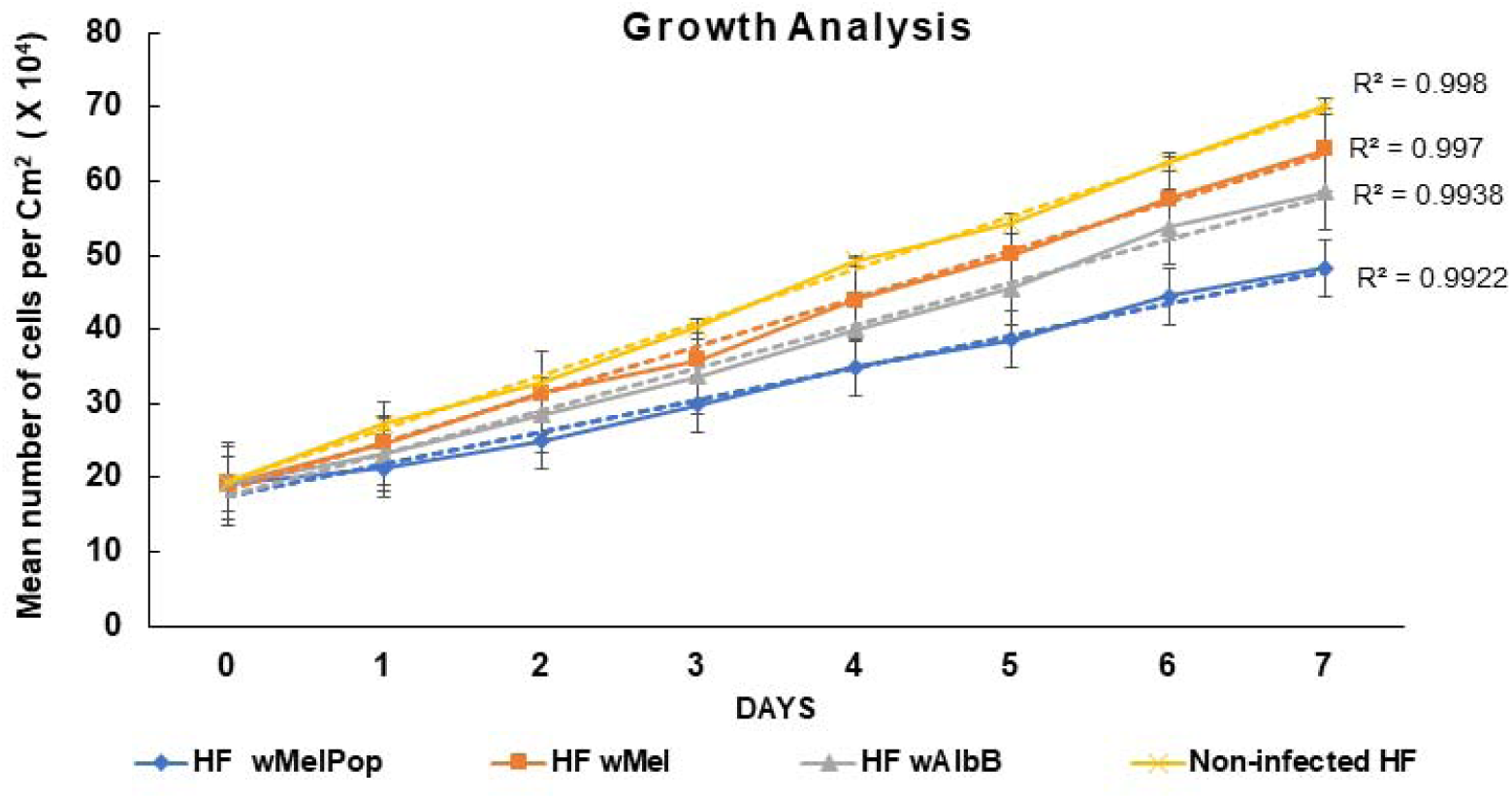
*In vitro* proliferation of the non-infected and differentially *Wolbachia* infected HIE-18 cell lines. Cells infected with *Wolbachia* (*w*AlbB, *w*Mel, and *w*MelPop) proliferated slowly, resulting in increased doubling time. Horizontal axis represents the mean number of cells per cm^2^ area + SD of three replicate culturing flasks.

### 3.6. Effect of culture media and temperature on *Wolbachia* growth in HIE-18 cells

*Wolbachia* infected HIE-18 cells in three different standard media (Schneider’s medium, Leibovitz’s L-15 medium, and Shield’s medium) and a mixture of Leibovitz’s L-15 and Schneider’s media (1:1) were grown for *Wolbachia* quantification after seven days of incubation. There was a significant effect of the Schneider’s medium on *w*AlbB strain density (Tukey test: *p*=0.02) and Schneider’s (Tukey test: *p*=0.01) and 1:1 mix of Schneider’s and L-15 media (Tukey test: *p*=0.03) on *w*MelPop density (Fig. 6) when compared to baseline. However, no effect of the media was seen on *w*Mel density.

**Fig. 6.**
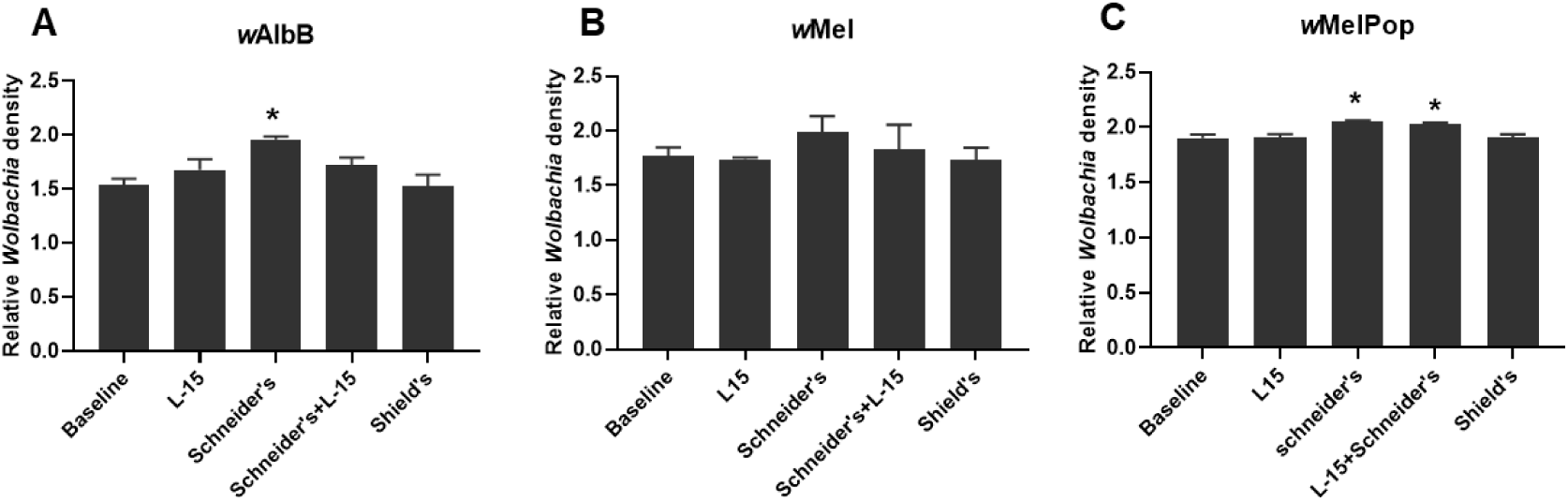
*Wolbachia* density in HIE-18 cells grown in different media. *Wolbachia* density was calculated relative to host *GAPDH* after growing infected HIE-18 cells in test medium for seven days. Here, baseline is *Wolbachia* density in HIE-18 cells grown in modified L-15 medium for 24 hours before assigning the test medium. **A.** *w*AlbB infected HIE-18 cell line. **B.** *w*Mel infected HIE-18 cell line. **C.** *w*MelPop infected HIE-18 cell line. Error bars are SEM calculated from three replicates of culturing flasks. Asterisks represent significance compared to baseline growth using Tukey’s test (\**p*<0.05).

Infected cells were grown at four different temperatures (26, 28, 30, and 32°C) and *Wolbachia* was quantified using a similar protocol to that described above (Fig. 7). Overall, seven days of incubation at assigned temperatures resulted in significant differences between temperatures in *Wolbachia* density of *w*AlbB-HIE-18 (one-way ANOVA: *F*_4,10_=22.97, *p*<0.0001), *w*Mel-HIE-18 (one-way ANOVA: *F*_4,10_=27.24, *p*<0.0001), and *w*MelPop-HIE-18 (one-way ANOVA: *F*_4,10_= 39.99, *p*<0.0001). However, there was a strain-specific response to temperature. In *w*AlbB-HIE-18 cells, *Wolbachia* density was highest at 28°C and decreased with increase in temperature above this (Fig. 7A). *w*Mel maintained highest density at 26°C (Fig. 7B), whereas *w*MelPop density increased with the temperature, highest at 30°C, but decreased at 32°C (Fig. 7C).

**Fig. 7.**
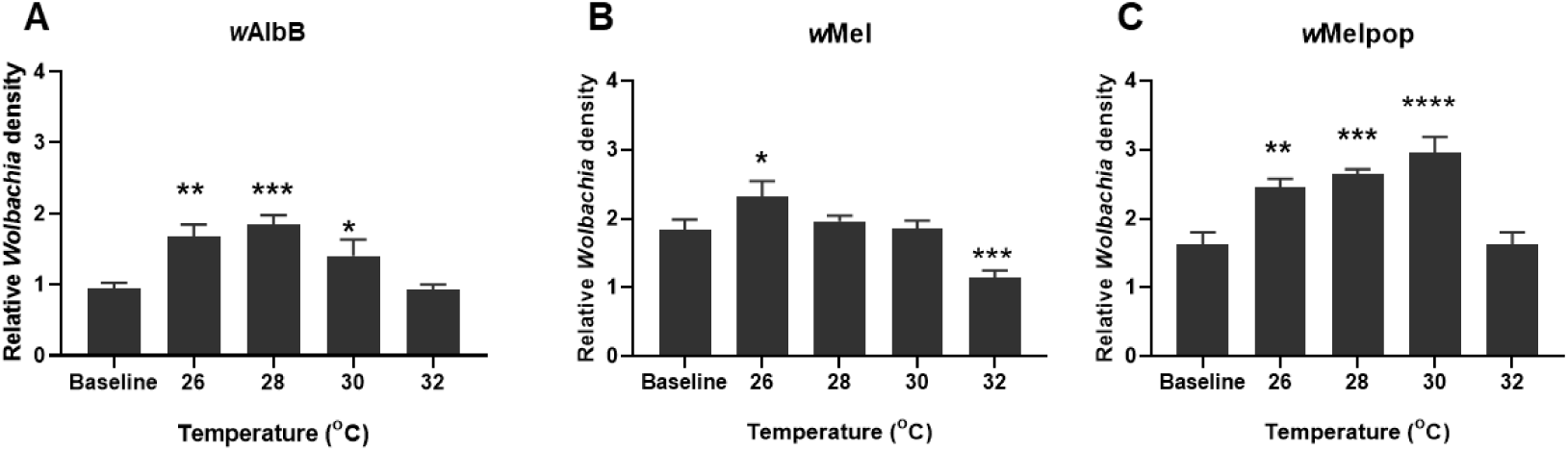
*Wolbachia* density in HIE-18 cells incubated at different temperatures. *Wolbachia* density is calculated relative to host GAPDH after growing infected HIE-18 cells at test temperatures for seven days. Here, baseline data is *Wolbachia* density in HIE-18 cells grown in modified L-15 medium for 24 hours at 30°C before assigning test temperature. **A.** *w*AlbB-infected HIE-18 cell line. **B.** *w*Mel-infected HIE-18 cell line. **C.** *w*MelPop-infected HIE-18 cell line. Error bars are SEM calculated from three replicate culturing flasks. Asterisks represent significance compared to baseline using Tukey’s test (\**p*<0.03, \*\**p*<0.005, \*\*\**p*<0.001, \*\*\*\**p*<0.0001).

### 3.7. Innate immune response of HIE-18 cell line against *Wolbachia*

Changes in *H. irritans* expression levels were investigated for the Imd (Immune deficiency) pathway transcription factor Relish (Fig. 8A-C), Toll pathway repressor Cactus (Fig. 8D-F), and three AMPs, Attacin, Cecropin, and Defensin (Fig. 8G-O) within 48 hours of initial infection with *Wolbachia* to better understand the *H. irritans* innate immune response towards the endosymbiont. There was a significant increase in Relish expression within 12 hours for *w*AlbB (Tukey test: *p*=0.005), *w*Mel (12 hours; Tukey test: *p*<0.0001), and *w*MelPop (12 hours; Tukey test: *p*<0.0001), which remained overexpressed for *w*Mel and *w*MelPop until 24 hours suggesting activation of the Imd signalling cascade, followed by deactivation. Cactus was significantly upregulated in comparison with the baseline at 12, 24 and 36 hours with a maximum at 24 hours (Tukey test: *p*<0.0001) but decreased after this time. AMPs are effector molecules for immune response and in HIE-18 cells all three AMPs tested were significantly upregulated within 36 hours of infection with few exceptions at 12 hours (see Fig. 8I, L, N). Overall, expression of AMPs was highest at 24 hours and decreased after this time.

**Fig. 8.**
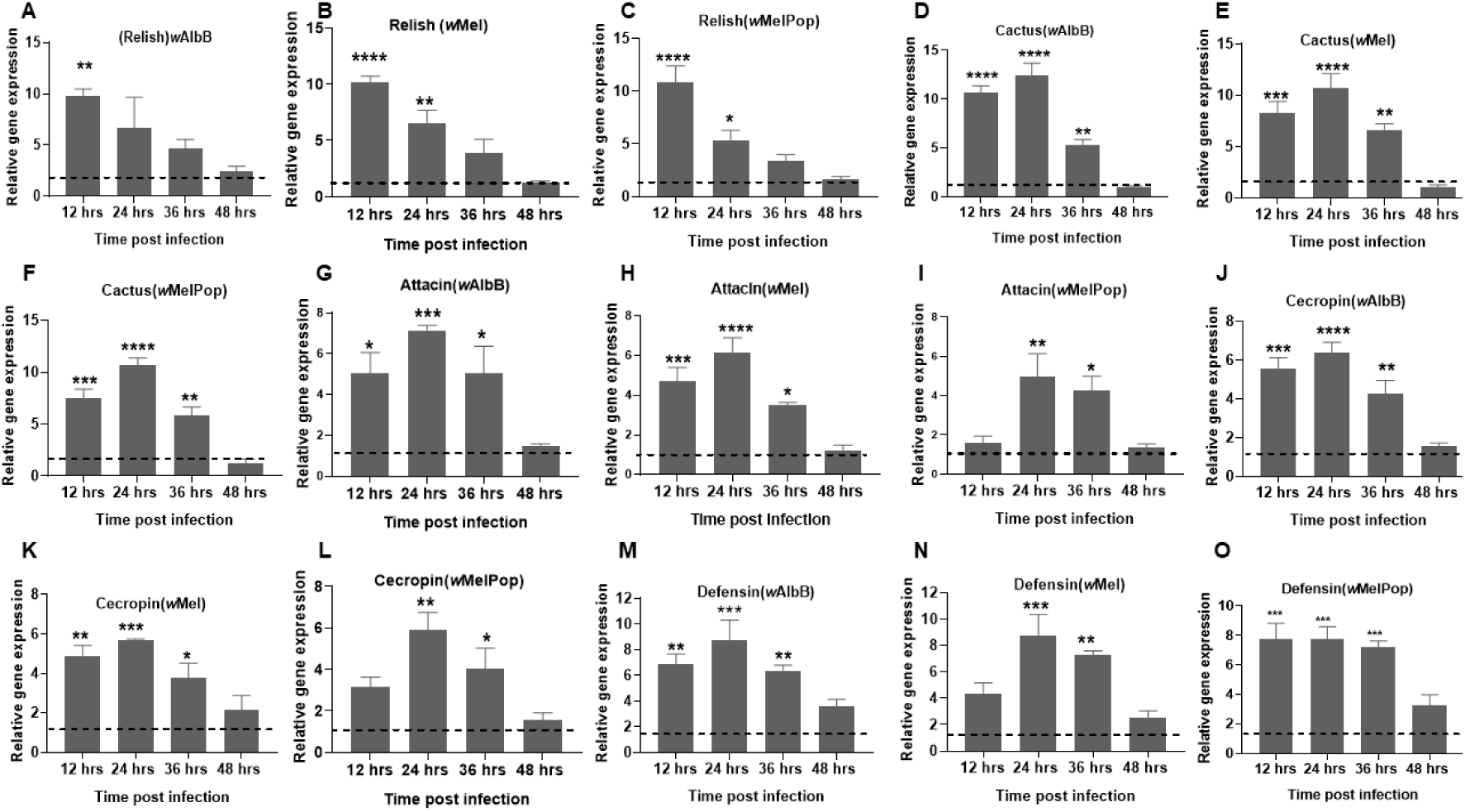
Relative gene expression of the immune pathways IMD, Toll, and AMPs in *H. irritans* determined by real-time PCR within 48 hours of infection with *Wolbachia* (*w*AlbB, *w*Mel, *w*MelPop). **(A-C)** Relative gene expression of *relish*, a regulator of the Imd pathways. **(D-F)** Relative expression of the *cactus* gene regulator of the Toll pathway. Relative gene expression of AMPs (G-I) *attacin* (J-L) *cecropin* (M-O) *defensin*. Error bar is SEM calculated from three independent experimental replicates. Quantification was normalised to the housekeeping gene *GAPDH,* and the dotted line here represents the relative gene expression of the non-infected control. Asterisks represent significance compared to control using a Tukey’s test (\**p*<0.03, \*\**p*<0.005, \*\*\**p*<0.001, \*\*\*\**p*<0.0001).

## 4. DISCUSSION

Insect cell lines play a critical role in understanding vital biological processes and insect-microbe interactions. Most mosquito cell lines were initiated with the intention of studying mosquito-arboviruses interactions ^49^. They have since been utilised in studying molecular interactions between hosts and endosymbionts that are unable to grow in the absence of host cells or tissues ^50–52^. Here we report the establishment of an *H. irritans* cell line (HIE-18) from embryos and studies of the interaction of HIE-18 cells with the endosymbiont *Wolbachia*.

Donor tissue plays a critical role in establishing insect cell lines. In the past, many attempts using specific adult tissues have resulted in limited success with relatively few significant exceptions ^53^. Currently, the use of whole embryonic and neonate larvae as an inoculant is a common approach as cells directly derived from this tissue often grow and proliferate better ^53^. We tried three different tissue sources as inoculum to initiate the primary cell line from HF: whole embryos (egg), neonate larvae and pupae. A major challenge to the establishment of the *Haematobia* cell culture from larvae and pupae was avoiding microbial contamination. Despite the use of multiple different disinfection protocols, fresh cattle manure used to maintain the source *Haematobia* colony was speculated to be a primary source of contaminant for the cell lines ^54^. In embryonic culture, one of the flasks HIE-18 with heterotypic cellular morphologies resulted in better cellular interaction leading to the successful establishment of a cell culture system. In the HIE-18 cell line, cell types with an approximately circular morphology have become predominant.

Frequently, primary cells have limited differentiation capabilities and undergo senescence within a short period (40-50 generations for mammalian primary cells) ^42, 55^. Sometimes, differentiation capabilities can be altered either naturally or through mutation resulting in a continuously proliferating cell system ^42^. The developed HIE-18 cell line has been subcultured through more than 200 passages, which equates to more than 400 generations suggesting that the HIE-18 cells have formed a continuous cell line. We have also thawed cryopreserved cells and in most instances the thawed cells have been found to be mitotically active and viable, indicating that HIE-18 cell line can be maintained for long term storage.

Horn flies are karyotypically diverse ^56^. Of three chromosomal investigations in HF two sourced insects from the USDA Livestock Insect Research Laboratory, Kerrrville TX, one from a wild population from Argentina and one from a population of HF from Uruguay ^56^. Horn flies from the USDA Lab colony had five pairs of chromosomes, without a sex chromosome ^57, 58^. However, the wild HF populations from Argentina and Uruguay had an extra B-chromosome and the resultant karyotype of 2n=11 (2n = 10+ B) in nearly 50% of the tested flies ^56^. Our cell line was established from embryos sourced from the USDA laboratory HF colony and had a similar chromosomal karyotype to that previously reported from the USDA flies ^58^.

Identification of appropriate media, and temperature for optimal cell growth, is a critical step in the establishment of a cell line. Growth conditions for early cell lines were usually determined by tedious experimentation and testing of different permutations of media components and growth conditions ^53^. The current preferred approach is to first test different commercially available media ^59^. Initially, the HIE-18 cell lines were established and cultured in modified Leibovitz’s L-15 medium designed for culturing a tick cell line by adding cholesterol, vitamins, glucose and amino acids to the L-15 medium ^44^. Based on our findings, we have recently moved to using Schneider’s medium for improved HIE-18 growth. Schneider’s medium is widely used for culturing *D. melanogaster* cell lines and differs in composition (mainly inorganic salts, vitamins, amino acids and cholesterol) from modified L-15 medium ^60^. Mostly, insect cells are cultured between 25 and 30°C and have been found to have reduced cell replications at lower temperatures ^61^. However, as a result of the study described here we are now maintaining the HIE-18 cells at 30°C for better proliferation.

Most available knowledge of insect cellular immune responses has been derived from studies with either *Drosophila* or mosquito cells, but the immune response in *Haematobia* species is less well understood. Cellular transinfection of insect cells with *Wolbachia* from different host backgrounds has shown mixed results in mosquitoes, and other dipteran species ^19, 34, 49^. It took a number of attempts to stably infect HIE-18 cell lines with *Wolbachia* as density in the infected HIE-18 lines decreased significantly in the first 48 hours after introduction. To better understand the initial interaction between the HIE-18 cells and *Wolbachia*, several gene expression studies were carried out with the Imd pathway transcription factor Relish, Toll pathway repressor gene Cactus, and three AMPs, Attacin, Cecropin, and Defensin. The immune gene expression study suggested that *w*AlbB, *w*Mel, *w*MelPop infections in the HIE-18 cell line activated the Imd pathway, resulting in the production of AMPs and reduced survival of *Wolbachia* until the infection became established in the HIE-18 lines. These results are consistent with the upregulation of the *Relish*, *Attacin*, *Cecropin*, and *Defensin* genes seen within 72 hours of infection of the LL-5 sand fly cell line with *w*Mel and *w*MelPop-CLA *Wolbachia*, and *Cactus* in the *Spodoptera frugiperda* Sf9 cell line infected with *w*Mel *Wolbachia* ^19, 34^. Despite the initial upregulation of immune response genes, HIE-18 cell lines have become stably infected with *w*AlbB, *w*Mel, and *w*MelPop which suggests a level of permissiveness of *Haematobia* species to *Wolbachia* or that *Wolbachia* is able to suppress or overcome the initial *H. irritans* innate response.

After the establishment of stable infections of *Wolbachia* in the HIE-18 cell line the optimal medium and temperature to achieve maximum densities of *Wolbachia* were determined. Microbial endosymbionts play an essential role in host metabolism by nutrition provisioning, and vice versa ^62^. Host nutrition and metabolism also influence endosymbiont dynamics ^62^. For example, supplementation with thiamine monophosphate decreased endosymbiont *Sodalis* and *Wigglesworthia* density in tsetse flies. Another similar example is supplementation of a *Drosophila* diet with sucrose which led to a higher density of *Wolbachia* in comparison to a yeast enriched diet ^62–64^. Temperature is another critical factor that can modulate host-endosymbiont interactions as it directly affects the rate of biochemical reactions, which can alter the performance and survival of organisms ^65^. Most temperature related studies have been done in either *Drosophila* or mosquito species and reported effects on *Wolbachia* density inside hosts can be complex and quite variable ^66–70^. Very little is known about the influence of culture medium and temperature in the context of different *Wolbachia* strain infected host cell lines. Interestingly, culture medium did not have any effect on *w*Mel density inside HIE-18 cells, but *w*AlbB and *w*MelPop had higher density in HIE-18 cells cultured in Schneider’s medium. *Wolbachia* strains differ in temperature preference as previously seen with other insect species ^68, 70^. This study has found that incubating *Wolbachia-*infected cell lines at 28°C will result in higher densities of *Wolbachia,* which will be required for future transinfection of *Haematobia* spp.

## 5. CONCLUSION

We have successfully developed a continuous cell line from *H. irritans* (HIE-18) and identified Schneider’s cell culture medium and 30°C incubation temperature as optimal for growth. The density of *Wolbachia* (*w*MelPop, *w*Mel, and *w*AlbB) decreased in the HIE-18 cells within 36 hours of infection, associated with the production of AMPs via the Imd immune pathway, but densities subsequently recovered. Currently, HIE-18 cells stably infected with these three strains of *Wolbachia* have been subcultured over 50 passages suggesting permissiveness of the HIE-18 cells for growth of this bacterium. The *Wolbachia* described here, adapted to and growing in high density in the *Haematobia* cell lines, will provide an important resource toward the development of novel control approaches for buffalo flies and horn flies.

## ACKNOWLEDGEMENTS

We thank Dr. Pia Olafson (USDA/ARS, Kerrville, TX) and Prof. Roger Moon (Entomology, Univ. MN) for making *H. irritans* eggs available for establishment of the *Haematobia* cell line. We thank Prof. Scott O’Neill (Monash University, Melbourne) and the Eliminate Dengue program for donation of the two *Wolbachia* strains *w*Mel and *w*MelPop used for this study. This project was funded by Meat and Livestock Australia.

